# The vomeronasal system decodes biological state from patterns of bile acid chemosignals

**DOI:** 10.64898/2026.04.30.721772

**Authors:** Varun Haran, Jinxin Wang, Mari Morimoto, Wen Mai Wong, Leena S. F. Rouyer, Jeffrey G. McDonald, Julian P. Meeks

**Affiliations:** Department of Neuroscience, University of Rochester School of Medicine and Dentistry, Rochester, NY, USA; NEUROCITY Program, City College of New York and University of Rochester, New York, USA; Graduate Program in Neuroscience, Department of Neuroscience, University of Texas Southwestern Medical Center, Dallas, TX, USA; Current affiliation: Salk Institute for Biological Studies, La Jolla, CA, USA; Center for Human Nutrition, Department of Molecular Genetics, University of Texas Southwestern Medical Center, Dallas, TX, USA

## Abstract

The rodent vomeronasal system, also known as the accessory olfactory system (AOS), detects excreted chemosignals that guide social, reproductive, and defensive behaviors. Excretions sensed by the AOS contain blends of organic chemicals that reflect the biological identity of the emitter, but it remains unclear how the AOS extracts useful information from these complex natural cues. Vomeronasal sensory neurons (VSNs) sense bile acids, a diverse class of molecules excreted by vertebrates in feces. Using mass spectrometry, we found that patterns of bile acid excretion – not unique molecules – provide sufficient information to decode the emitter’s species, diet, and gut microbiome status. Using population calcium imaging, we found that VSN populations demonstrate an inverse relationship between natural bile acid abundance and population response magnitude and support efficient information encoding through stimulus “whitening.” VSNs showed maximum sensitivity to taurine-conjugated bile acids, a novel class of vomeronasal ligands found at low natural abundance levels, that have high theoretical discriminatory value. VSN tuning patterns aligned strongly with theoretical axes for discriminating reptilian predators from vegetarians, and between mice with different gut microbiome states. These results reveal that the AOS extracts biological state from bile acid emission patterns through enhanced sensitivity to rare, information-rich ligands.

## Introduction

Chemosensation supports the development and maintenance of social behaviors across the animal kingdom (Buchinger, Li et al. 2014, Boesveldt and Parma 2021, Rokni and Ben-Shaul 2024). Despite decades of progress since the discovery of the peripheral mechanisms of mammalian chemosignal detection by olfactory receptors (ORs)(Buck and Axel 1991), taste receptors and channels (Adler, Hoon et al. 2000, Chaudhari, Landin et al. 2000, Nelson, Hoon et al. 2001, Tu, Cooper et al. 2018), and vomeronasal receptors (Dulac and Torello 2003, Keller, Baum et al. 2009, Suárez, García-González et al. 2012), several long-standing quandaries persist. Key among them is that we do not fully understand how combinations of naturally occurring chemosignal molecules are represented in the neural code, and how those neural codes relate to biologically significant parameters (*e.g.*, species, age, sex, health, etc.). This knowledge gap is especially pronounced in the mammalian vomeronasal system, also known as the accessory olfactory system (AOS), which begins in the vomeronasal organ (VNO).

The AOS has long been appreciated for its role in supporting physiological development and the expression of social, reproductive, and defensive behaviors (Stowers and Liberles 2016, Mohrhardt, Nagel et al. 2018), but several barriers have prevented mechanistic, systems-level investigations of the coding strategies used by neurons in the AOS. A primary barrier to progress has been a relatively limited understanding of the specific molecules detected by vomeronasal sensory neurons (VSNs) in the VNO. We and others identified several classes of non-volatile steroidal AOS chemosignals, including sulfated and other polar steroids (Nodari, Hsu et al. 2008, Haga-Yamanaka, Ma et al. 2014, Fu, Yan et al. 2015, Lee, Kume et al. 2019), and bile acids (Doyle, Dinser et al. 2016, Wong, Cao et al. 2020). Steroids and bile acids are excreted most prevalently through urine and feces, respectively (Doyle and Meeks 2018), and activate VSNs via Type 1 vomeronasal receptors (V1Rs) at low micromolar concentrations (Nodari, Hsu et al. 2008, Turaga and Holy 2012, Fu, Yan et al. 2015, Doyle, Dinser et al. 2016, Xu, Lee et al. 2016, Lee, Kume et al. 2019, Wong, Cao et al. 2020).

Steroids and bile acids have been studied extensively for their internal roles in development, sexual maturity, immune regulation, digestion, and metabolism (Choudhuri and Klaassen 2022), and there are many plausible biological benefits to animals that are capable of detecting these molecules in the environment. This has proven true in many vertebrate species, especially fishes (Buchinger, Li et al. 2014). However, naturally occurring excreted steroids and bile acids, with a few exceptions (Fu, Yan et al. 2015), are not exclusively associated with a specific biological state, but are instead components of complex blends (Nodari, Hsu et al. 2008, Ben-Shaul, Katz et al. 2010, Hofmann, Hagey et al. 2010, Haga-Yamanaka, Ma et al. 2014, Kaur, Ackels et al. 2014, Doyle, Dinser et al. 2016, Xu, Lee et al. 2016, Wong, Cao et al. 2020). This strongly suggests that the AOS, at least for steroidal molecules, may not be designed to interpret individual cues as behavioral triggers (*e.g.*, as canonical pheromones). Instead, the AOS may implement a combinatorial strategy, similar to the main olfactory system (MOS), to interpret steroidal chemosignal blends.

Here, we report a set of experiments exploring how VSNs encode the identity of chemosignal emitters based on patterns of bile acid information. We focus on bile acids because of their ubiquitous and well-studied biological function across vertebrate evolution (Hofmann, Hagey et al. 2010), and also because most known bile acids and conjugates are commercially available in pure form, facilitating their use as monomolecular stimuli. Using mass spectrometry, we identified patterns of bile acid excretion that varied across captive reptilian species (including mouse predators) and across mice with different gut microbiomes. We used *ex vivo* volumetric VSN Ca^2+^ imaging to determine whether, and how well, the AOS can use the available diversity of bile acid patterns to decode biological states. We found that VSNs perform stimulus “whitening” via an inverse relationship between natural bile acid abundance and population response magnitude. VSNs were especially sensitive to taurine-conjugated bile acids, a novel class of chemosignals present at low abundance levels in reptilian samples. VSNs detect taurine-conjugated bile acids with submicromolar sensitivity, and caused behavioral aversion in social and non-social contexts. Using multiple analytical models, we demonstrate that VSN bile acid tuning is aligned with theoretical axes supporting reptilian species discrimination, and strongly aligned with axes enabling discrimination among gut microbiome states. These results expand our understanding of bile acid detection by the AOS, and suggest that mouse vomeronasal periphery supports efficient encoding of biological state through stimulus whitening and alignment of neural sensitivities to naturally occurring bile acid emission patterns.

## Results

### VSNs can distinguish between reptilian species and diet using fecal cues

Previous research has documented the diversity of bile acid production and excretion across the vertebrate animal kingdom (Hagey, Vidal et al. 2010). The pattern of bile acid production and excretion has even been hypothesized to provide an “honest” fingerprint of the emitter’s identity (Hofmann, Hagey et al. 2010). Following conversations with Dr. Hagey, we established a collaboration with colleagues at the Dallas Zoo, which houses a large collection of reptilian species with known diets (including rodents, earthworms, insects, and vegetables, **Fig. 1A**). We prepared sterile-filtered aqueous extracts from fecal samples collected by the Dallas Zoo staff, and used them in a fluorescence Ca^2+^ imaging bioassay of VSN activity. This assay involves placing a freshly dissected vomeronasal epithelium in which VSNs express GCaMP6s into a custom chamber, facilitating simultaneous stimulus delivery to the epithelial surface and 3-dimensional time series imaging via objective-coupled planar illumination (OCPI) microscopy (**Fig. 1B**) (Holekamp, Turaga et al. 2008, Turaga and Holy 2012). The reptilian fecal extracts readily and reliably activated VSNs at 1000-fold dilution, and many VSNs showed activity patterns that were selective for individual samples (**Fig. 1C-E**).

**Figure 1.**
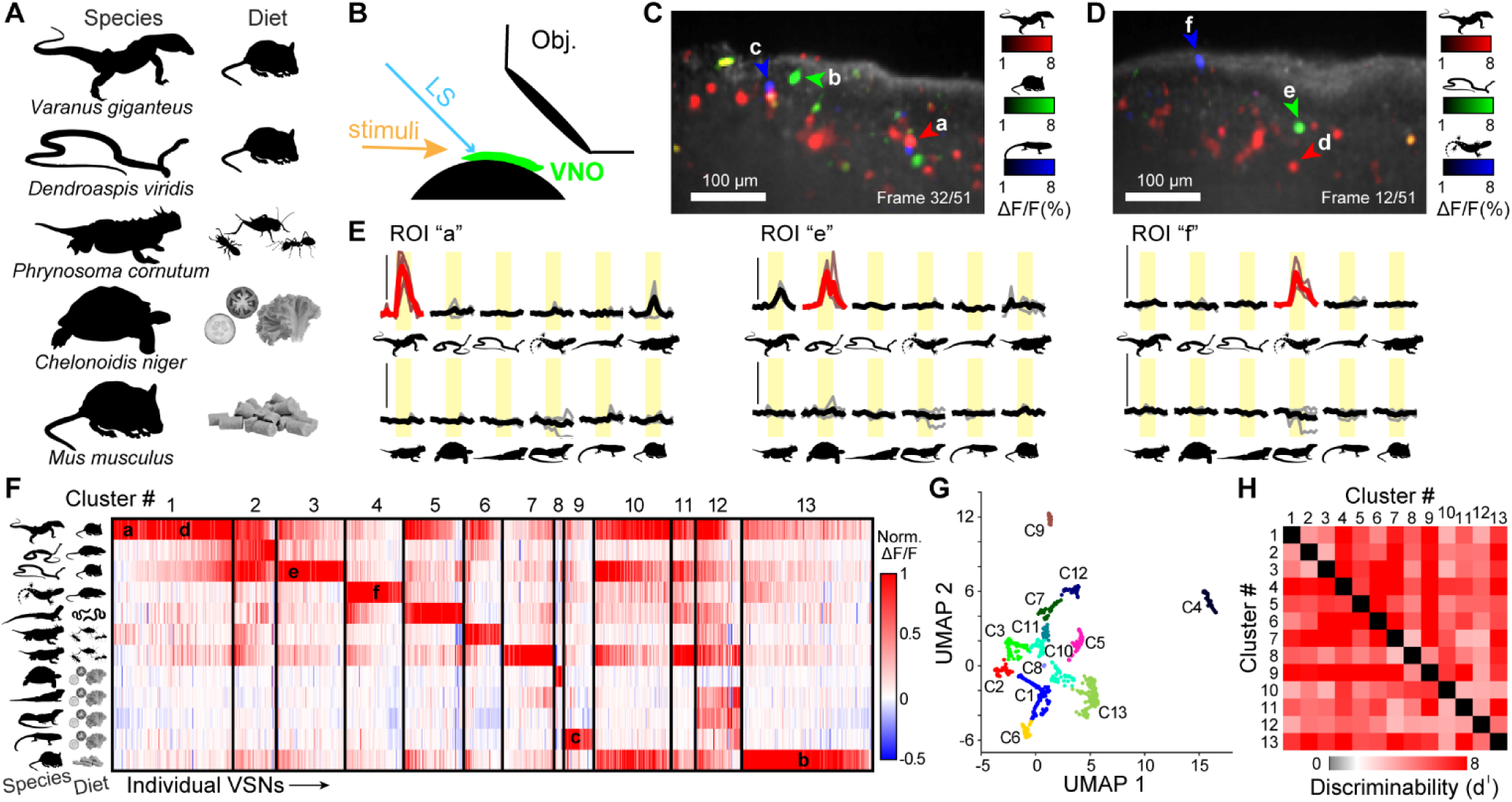
Fecal extracts from captive reptiles activate distinct populations of mouse vomeronasal sensory neurons (VSNs). (**A**) Highlights of the species and diets from which fecal samples were obtained. (**B**) Illustration of the experimental setup for objective-couple planar illumination microscopy of the vomeronasal organ (VNO) *ex vivo*. VSNs express GCaMP6s to enable live Ca^2+^ imaging. LS: light sheet. Obj.: imaging objective. (**C-D**) Individual frames from a 3-dimensional movie taken from a single VNO epithelium during stimulation with fecal extracts. Colored pixels reflect the mean across-trial GCaMP6s ΔF/F in response to stimulation to fecal extracts (diluted 1:1,000 in control saline). Arrowheads point to neurons highlighted in Panels E-F. (**C**) VSN responses to fecal extracts from *Varanus giganteus* (red), *Mus musculus* (green), and *Corucia zebrata* (blue). (**D**) VSN responses to fecal extracts from *Varanus giganteus* (red), *Dendroaspis viridis* (green), and *Helioderma horridum* (red). (**E**) Peri- stimulus time histograms of ΔF/F responses within regions of interest (ROIs) encompassing VSN somas. Thin traces reflect individual trials. Bold traces reflect across-trial mean. Yellow shading indicates the time of stimulation. Red colors indicate responses that were statistically significant (p<0.05, Student’s *t*-test) compared to control saline stimulation (not shown). (**F**) Clustered heatmap of normalized mean ΔF/F responses of >800 VSNs pooled across 5 mice. Each thin column reflects the response pattern of a single VSN all 12 stimuli. Vertical black bars indicate cluster divisions. (**G**) Uniform manifold approximation and projection (UMAP) plot of VSNs response patterns. Colored groups labeled C1-C13 refer to Clusters 1-13 in Panel F. (**H**) Heatmap of pairwise discriminability between the clusters in Panel G. High d^I^ values are associated with stronger statistical separation. White hue reflects the d^I^ equivalent of p = 0.05.

Evaluating the patterns of activation to the entire panel of 11 reptilian fecal extracts revealed the presence of multiple species-selective VSN populations (**Fig. 1F**). Several large populations of VSNs were selectively tuned to predator species, including species fed mice or rats in captivity, as well as species with invertebrate diets (**Fig. 1F**, Clusters 1-7). Two VSN populations demonstrated selectivity for vegetarian reptiles (Fig. 1F, Clusters 8-9). One sample, from the predator *Varanus giganteus* (Perentie monitor lizard) activated an especially large fraction of VSNs, including several that were activated by multiple reptilian species (Clusters 5, 6, 10, 11, and 12). VSNs in Clusters 1-9 rarely responded to dilute mouse fecal extracts, suggesting they detect cues enriched in reptiles. Overall, 511/807 (63.3%) of VSNs belonged to Clusters 1-9 with selectivity for reptilian fecal cues, 156/807 (19.3%) belonging to Clusters 10-12 demonstrating broad tuning, and 140/807 (17.4%) in Cluster 13, selective for mouse fecal chemosignals. Further analysis of these Clusters using dimensionality reduction and discriminability testing indicated that these Clusters were well-separated and statistically distinguishable. (**Fig. 1G-H**). The presence of these reptilian feces-selective VSN populations demonstrates that cues in fecal extracts can be used by the AOS to distinguish reptiles from mice, and reptiles from other reptiles.

### Patterns of bile acid excretion contain sufficient information to identify reptile species and diet

Prior molecular analysis of bile acid diversity across species have indicated that patterns of bile acid production and excretion vary extensively, including across reptilian species (Hagey, Vidal et al. 2010, Hofmann, Hagey et al. 2010, Mohanty, Mannochio-Russo et al. 2024). This observation indicates that bile acids could be used by the AOS to discriminate reptilian fecal cues from each other. To test whether the patterns of bile acid excretion in our collection of reptilian fecal extracts contain sufficient information to support their discrimination, we performed mass spectrometry analysis on these samples (Fig. 2). We specifically evaluated the abundance of 22 bile acids and conjugates (*e.g.*, taurine- or glycine-conjugated BAs) using high performance liquid chromatography followed by mass spectrometry (LC-MS; Fig. 2; see Methods). Due to technical limitations (*e.g.*, variability in the specific timing and handling of samples by zoo staff prior to cold storage, variable availability of deuterated standards, etc.), we were unable to determine the exact concentration of each bile acid, and therefore focused on the relative patterns of bile acid excretion (Fig. 2A-B). We found that the bile acids CA, ursocholic acid (UCA), ursodeoxycholic acid (UDCA), and DCA had the highest overall relative abundance across samples, whereas conjugated bile acids, especially taurine-conjugated bile acids, had low relative abundance (Fig. 2A). When relative abundance was normalized per molecule, it was evident that patterns of bile acid excretion varied extensively across these samples.

**Figure 2.**
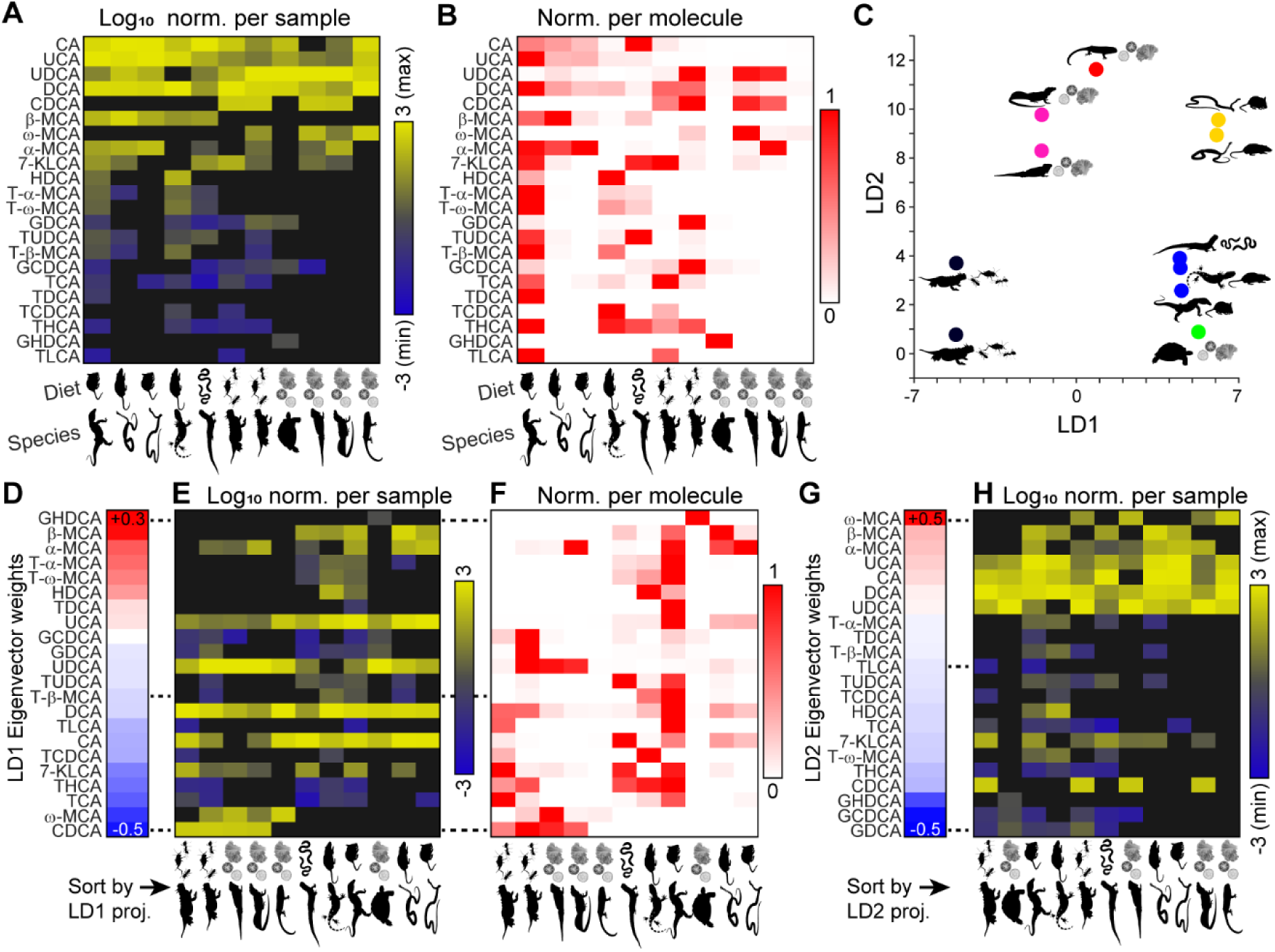
Mass spectrometry analysis of bile acid abundance in captive reptile feces across taxonomic suborder and diet. **(A)** Heatmap of log_10_-transformed abundance of 22 bile acids, normalized per sample to account for variable dehydration prior to collection by zoo staff. Each sample’s most abundant bile acid was multiplied by 10^3^ to make its log_10_ transform equal to 3. Undetectable molecules’ levels were set to 10^-5^ (displayed as black in the heatmap). **(B)** Heatmap of relative abundance, normalized by the maximum concentration of each bile acid across all samples. **(C)** Scatterplot displaying the projection of each sample’s bile acid abundance pattern (Panel A) along the first two linear discriminant eigenvectors (LD1, LD2). LD eigenvectors were calculated based sample taxonomic suborder and diet. **(D)** Heatmap of LD1 eigenvector weights, sorted from maximum-minimum. **(E, F)** Log10- transformed bile acid abundance (**E**, same as Panel A), and relative bile acid abundance (**F**, same as Panel B). The rows (bile acids) are arranged in the same order as LD1 eigenvector weights, and columns (samples) are arranged in the same order as LD1 projections (x-axis on Panel C). **(G, H)** Same as Panels D-E, but with bile acids (rows) arranged by LD2 eigenvector weights (**G**), and samples (columns) arranged by LD2 projection (y-axis on Panel C).

In order to determine whether and how bile acid content might be used to distinguish samples from one another, we used linear discriminant analysis (LDA; Fig. 2C). Rather than treat each sample as its own category, we grouped them based on their 4 taxonomic suborders (Autarchoglossa, Serpentes, Iguania, and Cryptodira), and 3 macroscopically similar diets (carnivore and vegetarian), generating 6 unique combinations of suborder and diet. LDA identified patterns of normalized bile acid abundance that could be used to discriminate between samples (Fig. 2C). Inspection of the primary LDA eigenvector (LD1) indicated which specific bile acid patterns supported optimal sample discrimination. For example, high normalized abundance of CA, UCA, alpha muricholic acid (α-MCA) and beta muricholic acid (β-MCA) supported discrimination of species in suborders Autarchoglossa and Serpentes that had rodent diets (Fig. 2D-F). Rodent-fed animals from suborder Autarchoglossa could be further distinguished from rodent-fed Serpentes based on higher relative abundance of several taurine-conjugated bile acids (Fig. 2F). Several other combinations of relative bile acid abundance were identified in LD1, and in LD2 (Fig. 2G-H). LDA based purely on animal suborder or diet were independently informative, but with weaker associations than the combination of suborder and diet (data not shown). Overall, these results support the hypothesis that reptilian fecal samples possess theoretically useful information for species and diet discrimination based on bile acid abundance patterns.

### VSN tuning to bile acids supports reptilian species discrimination via efficient encoding

After establishing that bile acid information in reptilian feces theoretically supports discrimination of different species and diets, we tested whether VSN bile acid tuning was aligned, or misaligned, with ideal discrimination axes (Fig. 3). We posited that if VSN tuning to bile acids was well-aligned with these axes, it would suggest an evolutionary/biological driver for AOS bile acid detection, for example to avoid reptile predators (Wang, Karbasi et al. 2023). To challenge this hypothesis, we selected 10 bile acids, representing a subset of the molecules assessed by mass spectrometry (Fig. 3A-B), as a stimulus panel for VSN Ca^2+^ imaging (Fig. 3C). We observed robust responses to all of the chosen molecules at 1 µM, a concentration at which selective responses to common bile acids were previously observed (Wong, Cao et al. 2020). Cluster analysis of the VSN responses to this stimulus panel indicated 20 functionally similar groups. Functionally separable VSN populations typically express the same vomeronasal receptor(s) (VRs)(Haga-Yamanaka, Ma et al. 2014, Lee, Kume et al. 2019, Wong, Cao et al. 2020), so this result indicates there are many more bile acid receptors exist than the 5 that have thus far been identified. Among the 20 clusters were several that responded to combinations of chenodeoxycholic acid (CDCA), 7-keto-lithocholic acid (7-KLCA), CA, and deoxycholic acid (DCA), molecules excreted at intermediate-to-high levels by several insectivore and vegetarian reptiles (Fig. 3C, Clusters 1, 2, 3, 6, 7, 12, and 17). A couple of rare VSN clusters responded to the glycine-conjugated bile acids glycodeoxycholic acid (GDCA) and glycochenodeoxycholic acid (GCDCA), which were observed in several species, including both carnivores and herbivores (Fig. 3B-C). We observed several large VSN clusters that responded to the taurine-conjugated bile acids taurocholic acid (TCA), taurochenodeoxycholic acid (TCDCA), and taurolithocholic acid (TLCA), which were excreted at low abundance levels by multiple carnivores, including mouse predators (Fig. 3C, Clusters 4, 5, 8-10, 12-14). Many of these VSNs also responded to 1 µM taurodeoxycholic acid (TDCA), a molecule detected at low abundance levels in Perentie monitor lizard feces (Fig. 3C, Clusters 15-16 and 19-20).

**Figure 3.**
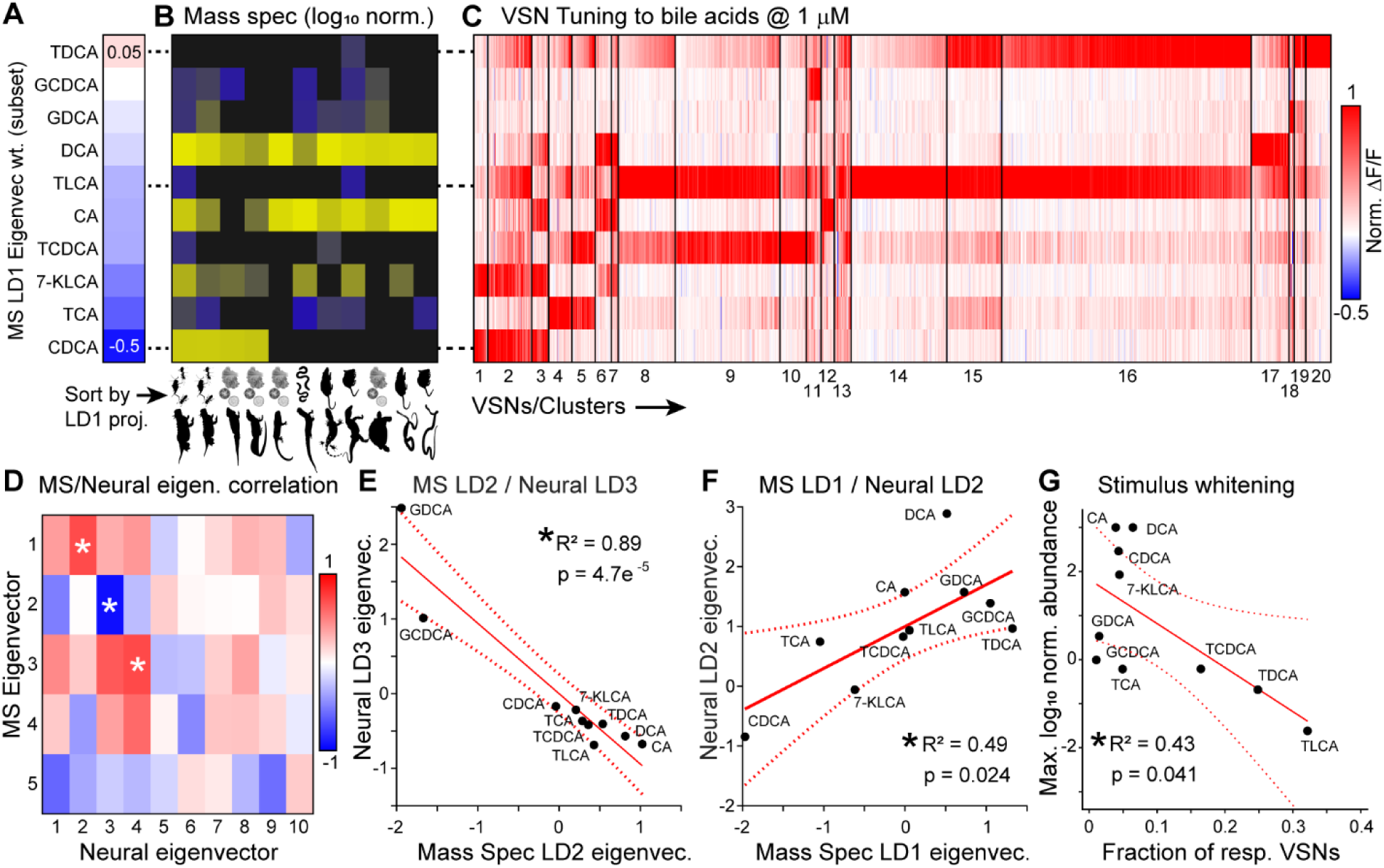
VSN bile acid tuning partially aligns with patterns that vary with reptile taxonomic suborder and diet. **(A)** Heatmap of mass spectrometry-based linear discriminant eigenvector (LD1) weights, sorted from maximum- minimum (subset of data from Fig. 2D). These 10 bile acids were chosen as stimuli for VSN tuning experiments. **(B)** Heatmap of log_10_-transformed abundance of the 10 bile acid ligands in Panel B (subset of data from Fig. 2E). **(C)** Clustered heatmap of normalized VSN responses to the bile acids highlighted in Panels A-B. All stimuli were delivered at 1 µM. Cluster divisions are denoted by vertical black lines. Clusters are arranged to visually highlight similarities with Panel B. **(D)** Heatmap of pairwise Pearson correlation “R” values between each mass spectrometry- based LD eigenvector and each neural response-based LD eigenvector. Asterisks denote R-values with associated p-values < 0.05. **(E)** Example correlation between the mass spectrometry-based LD2 eigenvector and neural response-based LD3 eigenvector. The ligands associated with each pair of weights is noted near its corresponding datapoint in the graph. **(F)** Same as in Panel E, but for mass spectrometry-based LD1 and neural response-based LD2. **(G)** Scatterplot comparing the fraction of responsive VSNs to the maximum log_10_ normalized abundance values from mass spectrometry. The anticorrelation indicates that at 1 µM, more VSNs respond strongly to bile acids with lower natural abundance than to bile acids with high natural abundance, indicating stimulus whitening.

The bile acid response patterns we observed in VSNs were partially aligned with patterns emitted by reptiles (Fig. 3A-F). We used linear discriminant analysis to identify eigenvectors (bile acid weight patterns) that best separated VSN clusters from one another (Fig. 3D-F). We performed correlation analysis on the eigenvectors from mass spectrometry (theoretical axes for discriminating reptile samples based on species and diet) and VSN Ca^2+^ imaging (axes separating VSN clusters from each other, Fig. 3D). The first neural linear discriminant eigenvector (LD1) did not correlate well with any of the 5 mass spectrometry eigenvectors, indicating that neural data were not optimally aligned with theoretical information available in bile acid patterns. However, neural eigenvectors LD2-LD4 correlated with mass spectrometry eigenvectors LD1-LD3, respectively (Fig. 3D-F). These instances of alignment between neural- and mass spectrometry-based eigenvectors indicate that VSN bile acid tuning produces information from feces that supports discrimination of reptiles based on species and diet.

We noted that VSN clusters that responded to 1 µM taurine-conjugated bile acids were more plentiful, had more VSNs per cluster, and included a high percentage of strong responses (Fig. 3C). Taurine-conjugated bile acids provided useful information for discriminating reptile species and diet, but were found at relatively low abundance levels (Fig. 3B). We compared maximal abundance of each bile acid across the 11 reptile samples to the fraction of response VSN population activity, finding a negative correlation (Fig. 3G). The increased amount of strong activity to taurine-conjugated bile acids at 1 µM suggested that VSNs are especially sensitive to this class of informative, but low-abundance chemosignals. The inverse relationship we observed between natural bile acid abundance and VSN population activity indicates that the AOS supports efficient encoding of reptile species and diet by balancing activity levels across log orders of natural abundance, a form of stimulus “whitening.”

### VSNs are highly sensitive to taurine-conjugated bile acids that cause natural aversion

Given the large number of 1 µM taurine-conjugated bile acid-responsive VSNs, we hypothesized that taurine-conjugated bile acids activate VSNs at very low concentrations. LCA and derivatives (e.g. TLCA) have previously been associated with acute cellular toxicity in some tissues (Graf, Kurz et al. 2002, Katona, Anant et al. 2009), which may partially account for their broad activation of VSNs. We therefore focused subsequent taurine-conjugated bile acid studies on TDCA, the second-most active AOS ligand identified in this screen. We conducted a concentration-response analysis for 3 bile acid ligands, DCA (0.01–10 µM), TDCA (0.001–1 µM), and GDCA (0.01–10 µM, Figure 4). We observed a range of sensitivities to these ligands, and identified groups of similarly-tuned VSNs using cluster analysis (Fig. 4A).

**Figure 4.**
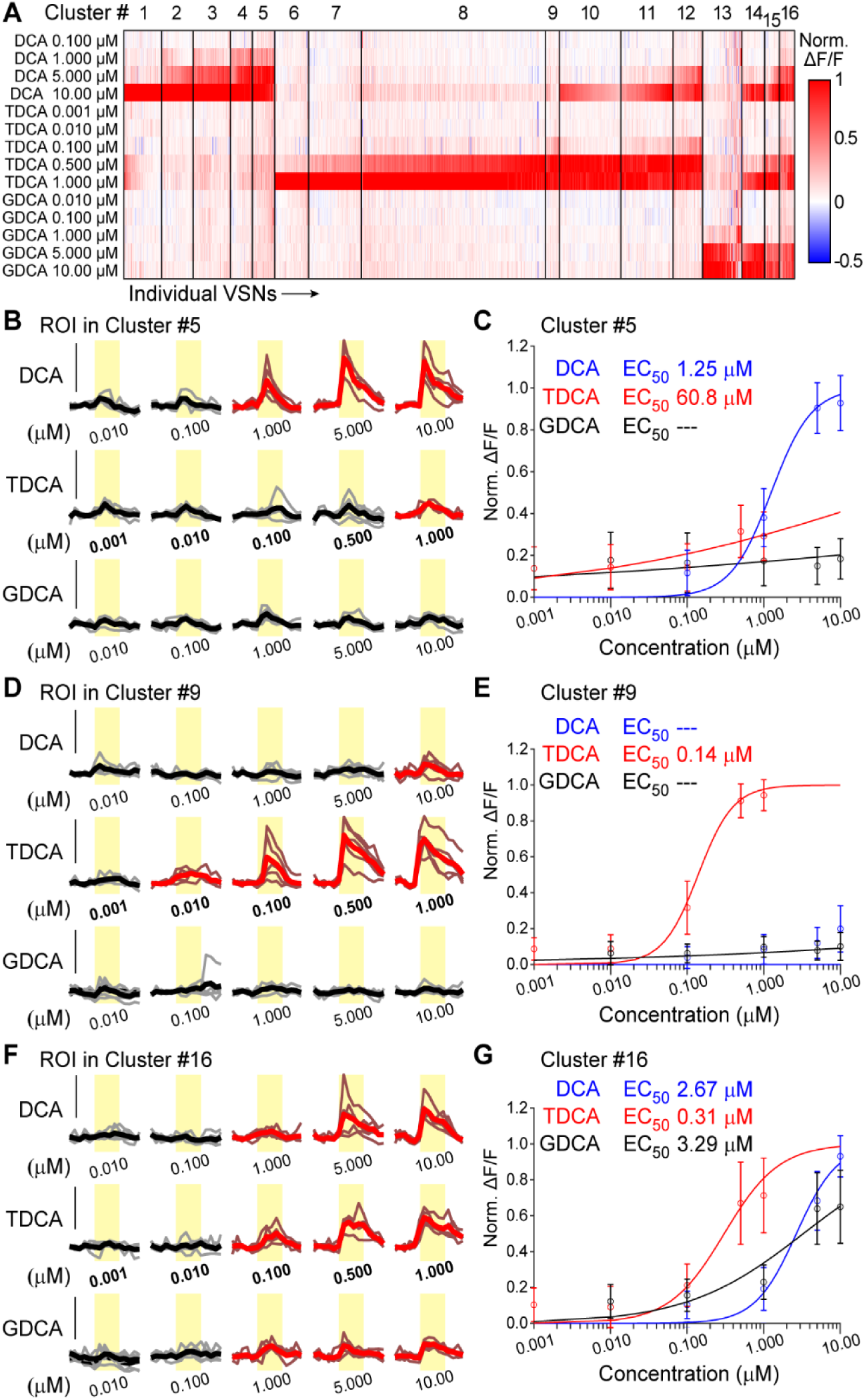
VSNs respond to taurine-conjugated bile acids at submicromolar concentrations. (**A**) Clustered heat map of 1,857 responsive VSNs exposed to DCA, TDCA, and GDCA at a range of concentrations (7 tissues from 2 males, 2 females). (**B, D, F**) Peri-stimulus time histograms of stimulus responses to ROIs from Cluster 5 (**B**), Cluster 9 (**D**), and Cluster 16 (**F**). Bold traces reflect mean responses to 5 stimulus repeats. Red traces are statistically larger than control (DCA 0.01 µM)(Wilcoxon Rank Sum Test, p<0.01). Yellow rectangles indicate the time of stimulus delivery. Vertical black bars indicate 40% ΔF/F. (**C**, **E**, **G**) Concentration-response curves from Cluster 5 (**C**), Cluster 9 (**E**), and Cluster 16 (**G**). Open symbols reflect mean Normalized ΔF/F signals across all cells in the cluster. Error bars reflect standard deviation of the mean. Solid lines reflect median solutions to the Hill Equation (applied to each cell in the cluster, see Methods). EC_50_ values for unresponsive ligands are noted as “- - -”.

This analysis identified VSN populations with a high degree of bile acid sensitivity and selectivity. Among these were 418 neurons (22.5% of analyzed cells) that were selective for DCA (Clusters 1-5, Fig. 4). Cluster 5 contained 62 VSNs (3.3% of analyzed cells) with high sensitivity and selectivity for DCA (Fig. 4B-C). VSNs in Cluster 5 had a median EC_50_ value for DCA of 1.25 µM (Fig. 4C). As in the initial screens (Figs. 1-2), many more VSNs responded to TDCA at 1 µM than the other ligands, despite being applied at a 10-fold lower concentration (Fig. 4A). Clusters 6-9 included VSNs that were selectively activated by TDCA (totaling 788 cells, 42.4%). Among these, the 39 VSNs (2.1% of analyzed cells) in Cluster 9 displayed the highest TDCA sensitivity (Fig. 4D-E). VSNs in Cluster 9 had a median EC_50_ value for TDCA of 140 nM (Fig. 4E). We also observed 396 VSNs (21.3% of analyzed cells) with high TDCA sensitivity that were also activated by DCA (Clusters 10-12, Fig. 4A). Among the 81 VSNs (4.4% of analyzed cells) in Cluster 12, the median EC_50_ for TDCA was 126 nM and the median EC_50_ for DCA was 5.64 µM. Cluster 13 contained 109 VSNs (5.9% of analyzed cells) with selective responsiveness to GDCA, with the median EC_50_ for GDCA of 2.17 µM. We observed 146 VSNs (7.9% of analyzed cells) that responded to DCA, TDCA, and GDCA (Clusters 14-16, Fig. 3A). Cluster 16, which contained 42 VSNs (2.3% of analyzed cells), displayed a median EC_50_ value to DCA of 2.67 µM, to TDCA of 310 nM, and to GDCA of 3.29 µM (Fig. 4F-G). These results confirm the taurine-conjugated bile acid potency on VSNs and suggest that TDCA and other taurine-conjugated bile acids, despite being present at lower concentrations in the environment, are highly active VSN ligands.

Given the presence of taurine-conjugated bile acids in rodent predator feces, we investigated their potential behavioral effects, using TDCA as a representative of the class (Fig. 5). We specifically tested whether TDCA, when added to natural blends of mouse fecal chemosignals, would be sufficient to drive behavioral changes. In each test, we studied both male and female mice and observed no sex-specific changes. We first performed a non-social two-choice assay using two cotton swabs dosed with female feces extracts alone, or female feces extracts spiked with TDCA (50 mg/ml, Fig. 5A-C). In this context, investigation time of the swab with spiked-in TDCA was markedly decreased, with no differences related to mouse sex (Fig. 5B-C). Next, we performed a two-choice assay in a semi-social context. Mice were maintained three per cage and fecal material was pooled for aqueous extraction. On the test day, two of the cage mates were anesthetized, painted with pooled fecal extracts with or without TDCA spike-in (50 mg/ml), and the unanesthetized cage-mate was allowed to freely investigate the animals for 10 min. We observed that mice spent more time investigating animals painted with their own fecal extracts without the TDCA spike-in, consistent with the TDCA effect in the non-social behavior context. These results indicate that TDCA, when added to a blend of familiar conspecific fecal cues, causes behavioral aversion. Furthermore, these effects reveal that modifying natural bile acid chemosignal patterns can alter their behavioral valence.

**Figure 5.**
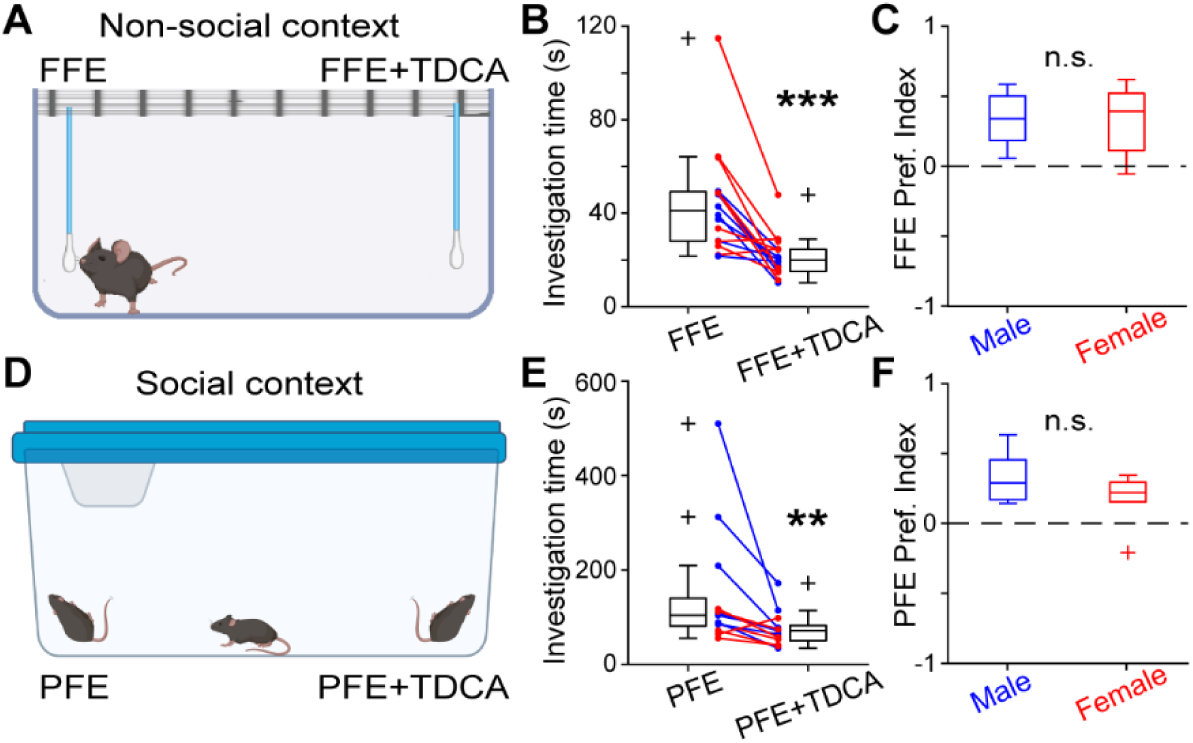
Spike-in of TDCA to fecal cues causes aversion in social and non-social contexts. **(A)** Non-social behavior setup. Mice were introduced to cotton swabs soaked with female feces extract (FFE) or FFE+TDCA for 10 minutes. **(B)** Investigation time in the non-social assay. Red symbols indicate female mice, blue indicate males. **(C)** FFE preference index for each sex. Positive values indicate preference for FFE over FFE+TDCA, vice-versa for negative values. **(D)** Social behavior setup. Groups of 3 mice were co-housed for a week, and pooled feces extracts (PFE) acquired. On test day, 2 of the 3 cage-mates were anesthetized and painted with PFE or PFE+TDCA (50 mg/mL). **(E)** Investigation time in the social assay. **(F)** PFE preference index for each sex. **p<0.01, ***p<0.001 (Wilcoxon signed rank test).

### VSNs demonstrate efficient encoding of gut microbiome states

The variability we observed in bile acid content across reptilian species adds to a large body of knowledge on this topic (Hagey, Vidal et al. 2010, Hofmann, Hagey et al. 2010). Prior work also indicates a strong contribution of the gut microbiome to bile acid emission (Wahlstrom, Sayin et al. 2016, Tang, Tang et al. 2023). To evaluate the impact of major alterations in microbiome status on bile acid emission, we collected feces from germ-free (gnotobiotic) and conventionally raised mice and measured bile acid abundance via mass spectrometry (Fig. 6). In this experiment, we were able to assess 59 detectable bile acid analytes, more than in the reptile samples, improving our depth of coverage (Fig. 6). Theoretical axes supporting discrimination of germ-free feces extracts (GF) from conventionally-raised feces (CF) extracts were identified using the same linear discriminant analysis used for reptilian samples, but with only 2 categorical variables (“GF” and “CF”), resulting in a single linear discriminant eigenvector (LD1) for bile acid abundance measurements (Fig. 6B). This analysis indicated very strong pattern separation between GF and CF samples, with GF samples being enriched in several taurine-conjugated bile acids, including tauro- α-muricholic acid (T-α-MCA), tauro-β-muricholic acid (T- β-MCA), tauro-ursodeoxycholic acid (TUDCA), TCA, and TCDCA. CF samples had low abundance levels of these same taurine-conjugated bile acids, but were instead enriched in other bile acids, including ω-muricholic acid (ω-MCA), ursocholic acid (UCA), 12-keto-lithocholic acid (12-KLCA), and CA (Fig. 6C). We chose these mouse microbiome-dependent bile acids as monomolecular ligands in a VSN Ca^2+^ imaging stimulus panel, again at the 1 μM concentration (Fig. 6C-E).

**Figure 6.**
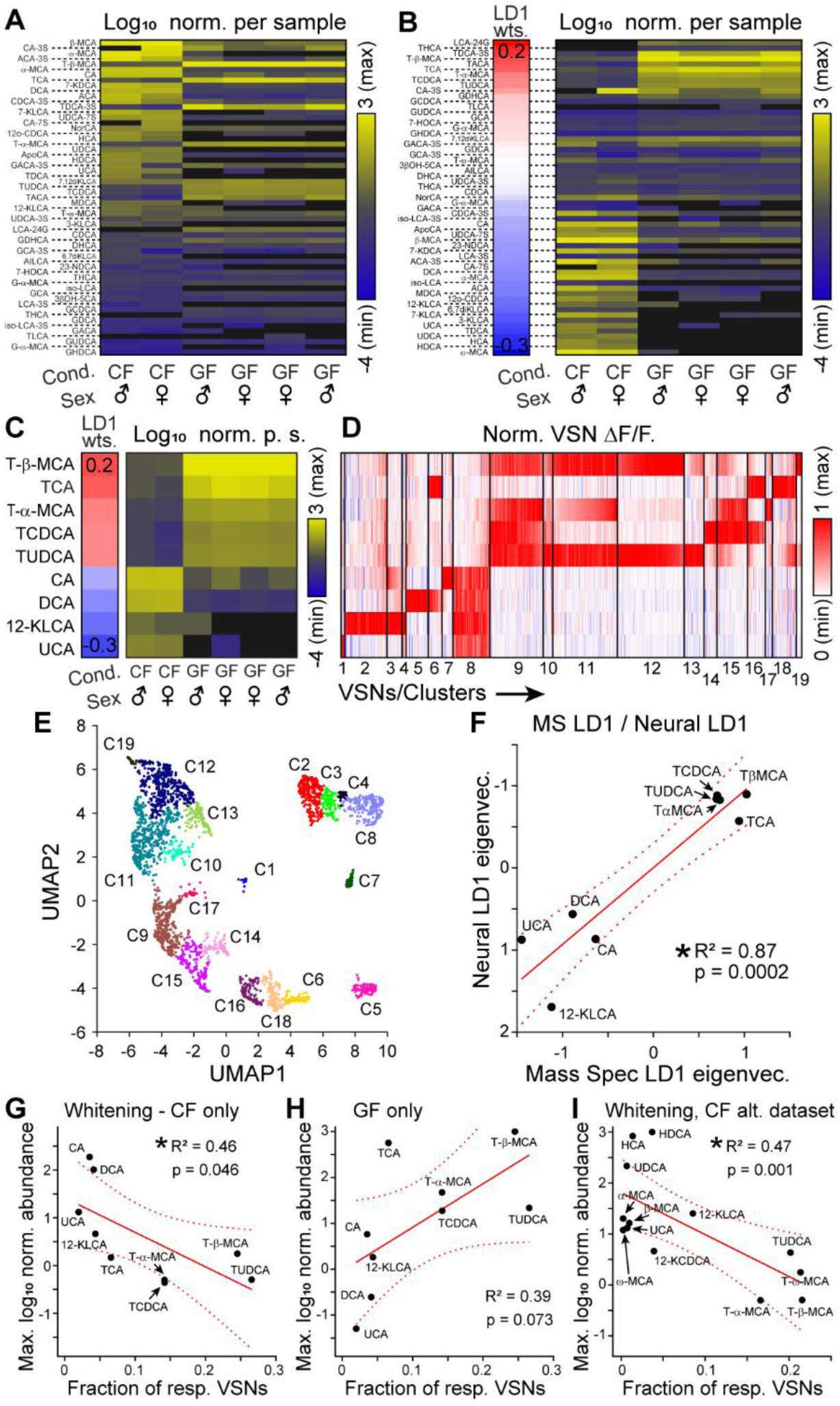
VSN tuning aligns with microbiome-dependent bile acid excretion patterns. **(A)** Heatmap of log_10_- transformed abundance of 59 bile acids, normalized per sample. Undetectable molecules’ levels were set to 10^-4^ (displayed as black in the heatmap). CF: conventionally raised feces extract, GF: germ-free mouse feces extract. **(B)** Mass spectrometry-based LD1 Eigenvector weights (left) and reorganized MS data (from Panel A). Rows match LD1, and columns are organized by projection of samples along MS LD1. **(C)** Subset of data in (B), with bile acid ligands matching VSN tuning data in (D). **(D)** Heatmap of VSN tuning to bile acids shown in (C), with rows organized as in (C) and columns arranged based on approximate alignment with MS LD1. **(E)** UMAP projections of VSN clusters in (D). Note the strong separation between clusters aligned with conventionally raised mouse feces (“CF”)(Clusters C1-8) and germ-free mouse feces (“GF”)(Clusters 9-19). (**F**) Comparison of LD1 weights based on mass spectrometry (x-axis) and LD1 weights based on VSN tuning (y-axis). Each dot represents an individual bile acid’s projection along each LD eigenvector. The solid red line indicates the slope of the linear regression best fit, and dashed lines the 95% confidence intervals of the linear fit. (**G**-**H**) Scatterplot comparing the fraction of responsive VSNs to the maximum log_10_ normalized abundance values from CF samples only (**G**) or GF samples only (**H**). (**I**) Same as G (CF-only abundance values), with VSN responses taken from an alternate stimulus set.

Cluster analysis identified VSN populations that responded to bile acids strongly associated with CF samples, but not to bile acids associated with GF samples (Fig. 6D, Clusters 1-8), and vice versa (Fig. 6D, Clusters 9-19). The exception was Cluster 6, which responded to both DCA (associated with CF samples) and TCA (associated with GF samples). Dimensionality reduction methods highlighted the separation between Clusters 1-8 and 9-19, with Cluster 6 being intermediate (Fig. 6E). Comparing the linear discriminant eigenvector (“Mass Spec LD1”) to the primary neural equivalent (“Neural LD1”) revealed a strong correlation (Pearson R^2^ = 0.87, p = 0.002), indicating VSN tuning extracts most of the available information about gut microbiome state present in bile acid emission patterns (Fig. 6F).

Given the evidence that VSNs support efficient encoding of naturally occurring patterns of bile acid emission in reptile samples (Fig. 3), we hypothesized that VSNs would show similar evidence of efficient encoding to naturally occurring conspecific bile acid patterns. Conversely, we hypothesized that this inverse relationship would not be observed for bile acid emission patterns from germ-free mice, which are intentionally unnatural. VSN responsivities to bile acid abundance levels from CF samples indeed showed an inverse correlation (Pearson R^2^ = 0.46, p = 0.046; Fig. 6G), whereas no such relationship was observed for bile acid abundance patterns in the “unnatural” GF samples (Pearson R^2^ = 0.39, p = 0.073; Fig. 6H). In fact, this analysis indicated that VSN activation showed a trend towards a positive correlation for bile acids emitted by germ-free mice, inconsistent with efficient encoding of unnatural bile acid emission patterns. Additional tests of VSN tuning to an expanded panel of bile acids further demonstrated the strong inverse correlation between neural tuning and CF bile acid abundance levels (Pearson R^2^ = 0.47, p = 0.001; Fig. 6I).

### VSN bile acid tuning encodes a combinatorial representation of gut microbiome state

The alignment we observed between mass spectrometry information and bile acid tuning (Figs. 3 and 6) supports the theoretical capacity for VSN bile acid tuning to discriminate between biologically relevant states. However, these analyses did not directly determine whether VSN bile acid response patterns match sensitivities to fecal extracts themselves. We therefore included dilute GF and CF extracts, taurine-conjugated bile acids, and unconjugated bile acids in a combined VSN stimulus panel (Fig. 7). Cluster analysis of VSN tuning patterns to this panel were highly consistent with predictions based on mass spectrometry and monomolecular tuning analysis alone (Fig. 7A). Specifically, we identified VSN clusters that were activated by 1:300 diluted male and female CF extracts, but not 1:300 diluted male and female GF extracts (Fig. 7A, Clusters 1-3). Cluster 3, the largest of these clusters, comprised VSNs that selectively responded to 1:300 CF extracts, and also responded to 1 μM CA and 1 μM DCA, which were upregulated in CF samples (Fig. 7A, also see Fig. 6C-D). We measured the percentage of VSNs with a statistically significant ΔF/F response to both CF extracts and bile acid ligands, which confirmed the association of CF-responsive VSNs with CA and DCA responsivity (Fig. 7B). Other VSN clusters in this assay were sensitive to 1:300 diluted male and female GF extracts, but not to CF extracts (Fig. 7A, Clusters 11, 16-22). All but one of these clusters (Cluster 22) was responsive to at least one taurine-conjugated bile acid shown to be enriched in GF extracts (Fig. 7A). We measured the percentage of VSNs with overlapping responsiveness to both GF and monomolecular bile acids, confirming that the VSNs responsive to GF were also responsive to bile acids identified as enriched by mass spectrometry (Fig. 7C).

**Figure 7.**
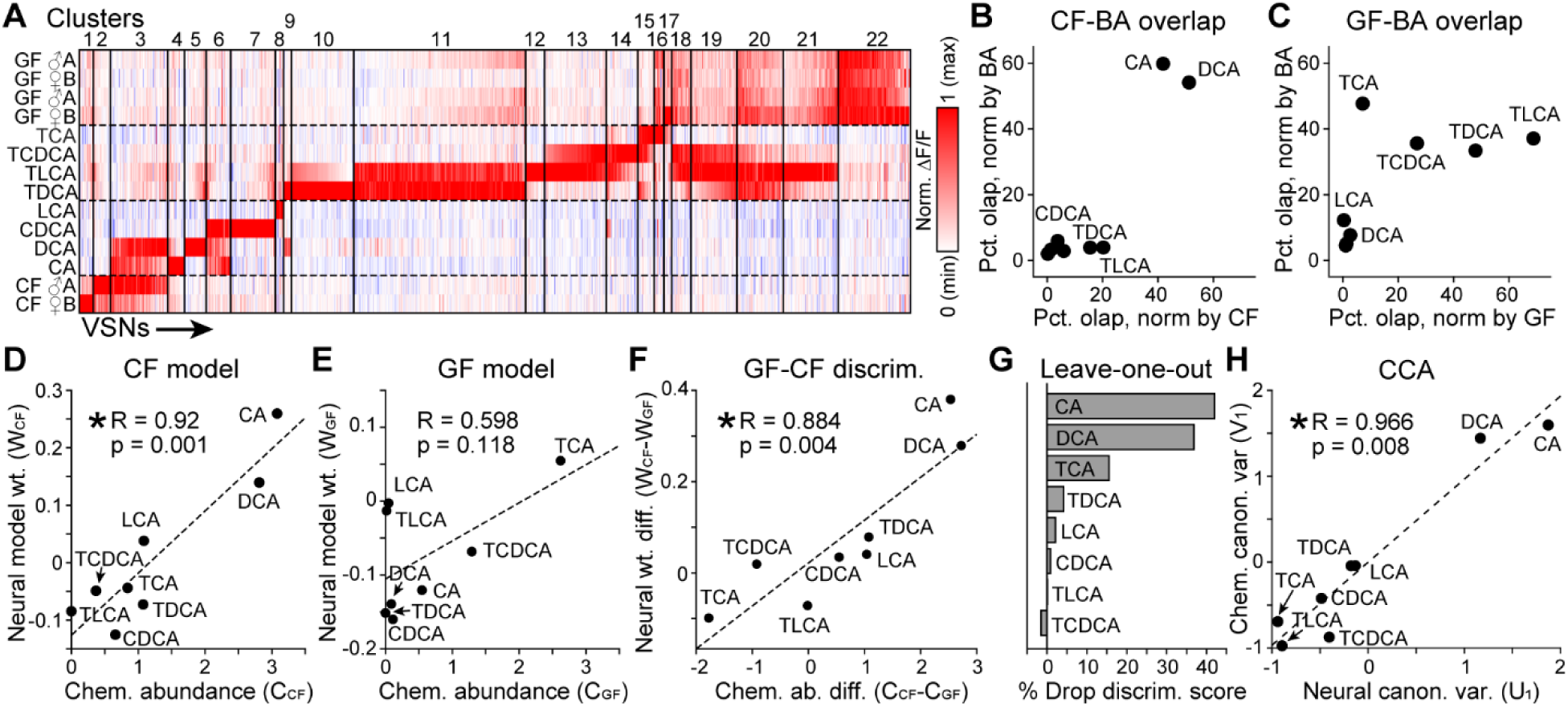
VSNs distinguish germ-free feces (GF) from conventionally-raised feces (CF) using bile acid information. **(A)** Clustered heatmap normalized VSN responses to GF extracts, CF extracts, taurine-conjugated bile acids, and unconjugated bile acids. Data from 5,117 VSNs were pooled across experiments on 12 VNOs from 8 mice (3 males, 5 females). Vertical pixels reflect individual VSNs, vertical black bars separate clusters. Dotted lines visually separate groups of similar stimuli. **(B-C)** Comparison of VSN response overlap between CF extracts **(B)**, GF extracts **(C)**, and each bile acid stimulus. The total number of VSNs with overlapping responsivity in each group was normalized by the total number of CF-responsive VSNs (x-axis) and by the total number of VSNs responding to each bile acid. (**D-E**) Correlation analysis between bile acid abundance (x-axis) and neural model weights (y- axis) indicates strong correlations for CF extracts **(D)**, but weak correlation for GF extracts **(E)**, evaluated independently. **(F)** Comparison of bile acid abundance differences in CF and GF extracts to neural weight differences. **(G)** Comparison of the amount of reduction in discrimination score (% Drop discrim. score, x-axis) by removing each bile acid from the model via leave-one-out analysis. **(H)** Canonical correlation analysis of neural (x- axis) and chemical (y-axis) canonical variates reveals a strong correlation between bile acid tuning and bile acid variability across GF and CF extracts.

Cluster analysis (Fig. 7A–C) indicated clear qualitative overlap between VSN responses to pure bile acids and fecal extracts. We proceeded to determine whether population-level tuning to these individual molecules quantitatively predicts the neural representation of microbiome state. We applied a multiple linear regression model mapping the response patterns of more than 5,000 VSNs onto their responses to dilute CF and GF extracts (Fig. 7D–H). Neural encoding weights for the CF extract (W_CF_) were highly correlated with the actual chemical abundance of these bile acids in CF feces (R = 0.920, p = 0.001; Fig. 7D). In contrast, the correlation for the GF state was weaker and did not reach significance (R = 0.598, p = 0.118; Fig. 7E), consistent with the reduced bile acid diversity in the absence of microbial metabolism. Discrimination between microbiome states, however, depends not on how faithfully VSNs encode each state in isolation, but on how their differential responses map onto chemical differences between states. We therefore compared the shift in neural encoding weights (ΔW = W_CF_ − W_GF_) against the shift in chemical abundance (ΔC = C_CF_ − C_GF_). This discrimination axis yielded a strong correlation (Pearson R = 0.884, p = 0.004; Spearman ρ = 0.952; Fig. 7F), demonstrating that vomeronasal sensitivity mirrors the chemical abundance shifts in the absence of gut microbiota. To determine which molecules drive this population-level discrimination, we performed a virtual knockout analysis (Fig. 7G). The unconjugated bile acids CA and DCA were the dominant drivers, accounting for 47.5% and 37.5% of the model’s discrimination score, respectively. The conjugated bile acid TCA provided the strongest opposing contribution (8.7% of the discrimination score), reflecting its enrichment in the GF state. We further validated this integrated chemical–neural relationship using canonical correlation analysis (CCA), which confirmed a highly significant multidimensional alignment between the neural and chemical data (canonical R = 0.966, p = 0.008; Fig. 7H). Collectively, these analyses reveal that VSN bile acid tuning constitutes a combinatorial neural code: one in which sensitivity to each molecule proportionally tracks its microbiome-dependent abundance shift, enabling population-level decoding of conspecific gut microbiome status.

## Discussion

The AOS has long been recognized for its role in detecting socially relevant chemosignals, yet the principles by which it encodes complex, naturally varying chemical blends have remained poorly understood. The mouse AOS, we show, discriminates biologically relevant states – including reptilian species, dietary categories, and conspecific gut microbiome status – through population-level encoding of fecal bile acid patterns. By combining chemical, physiological, behavioral, and computational approaches, we show that the AOS functions as a combinatorial decoder of fecal metabolic states, extracting biologically meaningful information from complex bile acid blends. While VSN bile acid tuning partially supports the discrimination of reptilian predators, its population-level encoding is very well aligned with the dynamic chemical output of the conspecific gut microbiome. These findings reframe the AOS not as a system of parallel single-molecule “labeled lines,” but as one that employs a combinatorial code to evaluate physiological and metabolic status of other animals.

### Decoding species, diet, and predator identity

VSN populations are selectively activated by reptile fecal extracts, demonstrating that the AOS can use these complex cues to distinguish reptiles from mice and to discriminate among multiple reptilian species. This extends earlier work showing that feces are a robust source of AOS ligands and that VSNs can distinguish bile acid signatures associated with sex and species in mammals (Doyle, Dinser et al. 2016, Wong, Cao et al. 2020). Linear discriminant analysis of fecal profiles revealed that normalized bile acid abundance contains sufficient information to separate samples by reptilian taxonomic suborder and coarse diet class (*e.g.*, carnivore, vegetarian), indicating that a neural decoder with access to the full bile acid profile could recover both species-level and ecological information. In an ecological context, such selective responses are consistent with a role for the VNO in monitoring the presence and identity of potential predators or competitors, functioning in parallel with known circuits that encode predator imminence and drive defensive behaviors (Papes, Logan et al. 2010, Wang, Karbasi et al. 2023, Nguyen, Rocha et al. 2024).

The array of water-soluble molecules present in fecal samples is immense, but prior work indicated that bile acids were among the most abundant polar molecules in aqueous extracts (Doyle, Dinser et al. 2016). We focused our work on bile acids in reptilian samples because others had hypothesized that observed changes in bile acid composition across vertebrate species and diets may serve as molecular “fingerprints” of biologically relevant qualities (Hagey, Vidal et al. 2010, Hofmann, Hagey et al. 2010). Across our extensive tests of monomolecular bile acid stimuli on the VNO, it was clear that VSN populations tile this chemosensory space densely. Even so, these studies have probed just a small fraction of the potential AOS ligands present in fecal samples. Future studies will be necessary to identify and validate other classes of fecal chemosignals engaging the AOS.

The presence of dozens of new natural AOS bile acid ligands activating functionally distinct VSN populations suggests bile acid chemosensation is far more extensive than previously appreciated. Prior work identified five vomeronasal receptors responsive to four unconjugated bile acids (Wong, Cao et al. 2020). Other studies have identified strong linkage between patterns of chemosensory sensitivity and vomeronasal receptor expression (Isogai, Si et al. 2011, Haga-Yamanaka, Ma et al. 2014, Kaur, Ackels et al. 2014, Isogai, Wu et al. 2018, Osakada, Ishii et al. 2018, Lee, Kume et al. 2019). This suggests that many – likely dozens or more – undiscovered bile-acid-responsive vomeronasal receptors exist. If this holds true, bile acid molecules and their chemosensory receptors will represent a large fraction of the known natural ligands of the mouse AOS. The receptors for most newly-discovered ligands remain to be discovered, but are likely members of the vomeronasal type 1 receptor (V1R) family, which are sensitive to bile acids and other polar sterols (Isogai, Si et al. 2011, Haga-Yamanaka, Ma et al. 2014, Lee, Kume et al. 2019, Wong, Cao et al. 2020). Importantly, our results indicate these to-be-determined receptors endow VSNs with the capacity to provide the brain with a combinatorial substrate for discriminating across biological states based on patterns of bile acid abundance.

### Enhanced VSN sensitivity to rare chemosignals with aversive qualities

One of the most striking results from this study was the inverse relationship between natural bile acid abundance and vomeronasal sensitivity (Fig. 3G). VSN population responses to 1 μM bile acids with high abundance across samples (*e.g.*, CA and DCA) was much lower than to bile acids with low abundance (*e.g.* TDCA, TCDCA; Fig. 3G). This increased responsiveness was supported by VSN populations that exhibited submicromolar sensitivity to taurine-conjugated bile acids, including VSNs displaying median EC_50_ values in the 100 nM range, representing an approximately 10-fold increase in sensitivity relative to unconjugated counterparts. The inverse relationship between environmental abundance and neural sensitivity produces – at the level of peripheral receptivity – a component of stimulus “whitening,” balancing the levels of neural activity generated by stimulus qualities across orders of magnitude of natural abundance (Friedrich and Wanner 2021, Chapochnikov, Pehlevan et al. 2023, Compton, Roop et al. 2023). This inverse relationship between natural bile acid abundance and VSN population activity was also observed in conventionally raised mouse feces, but not for germ-free mouse feces (Fig. 6). This suggests that this inverse relationship, which supports efficient encoding, is not present for any observable bile acid emission pattern, but is instead specific to abundance patterns most likely to be encountered in nature. This result is also noteworthy because the AOS achieves this type of stimulus decomposition in the periphery, likely through the evolution and maintenance of specialized chemosensory receptors, rather than in the brain via specialized neural circuitry.

The specific enrichment of taurine-conjugated bile acid sensitivity intersects aspects of the hepato-biliary circulation, through which bile acids are generated, modified, and recycled in a gut-microbiome-sensitive manner (Chen, Liu et al. 2026). In homeostatic conditions, these bile acids are likely to be rare in the environment, and their presence may signify the presence of a foreign species (*e.g.* a reptilian predator), or a conspecific with abnormal gut function, such as a sick or starving animal. Consistent with this interpretation, adding TDCA to familiar, conventional fecal extracts was sufficient to drive behavioral aversion in both social and non-social contexts, with no sex-specific differences (Fig. 5). This result demonstrates that a single, low-abundance bile acid class predictably alters behavioral responses to a familiar chemosignal blend. The AOS’ heightened sensitivity to taurine-conjugated bile acids may act as a generalized warning signal to drive avoidance of potentially dangerous biological states, whether they originate from predators or dysbiotic conspecifics.

### A combinatorial code for assessing gut microbiome status

Our comparison of chemical and neural coding strategies indicates that VSN bile acid tuning is partially aligned with the chemical patterns that distinguish reptilian species and diets, tuning was strongly aligned with patterns distinguishing conspecific gut microbiome states. Across multiple independent analyses – encoding models, discrimination axes, virtual knockouts, and canonical correlation analysis – we observed strong, statistically significant correspondence between VSN sensitivity profiles and microbiome-state-dependent bile acid variability. The degree to which each bile acid drives differential VSN responses between conventional and germ-free extracts was proportional to the magnitude of its differential abundance in these samples. This establishes a quantitative mapping in which vomeronasal sensitivity mirrors the structure of microbially driven chemical variation. The neural discrimination axis did not rely on a single biomarker but rather proportionally tracked the specific chemical abundance shifts induced by the microbiota. This distribution reflects the enzymology of microbial bile acid transformation: an intact microbiome converts primary bile acids into secondary forms and deconjugates taurine-conjugated species (Chen, Liu et al. 2026). Because the gut microbiome is highly sensitive to diet, infection, stress, and age, this combinatorial tuning provides mice with a continuous readout of the physiological fidelity of their social partners.

These results raise the broader question of why the mouse AOS might evolve such refined capacity to evaluate conspecific gut microbiome status. The gut microbiome profoundly influences host physiology, including immune function, metabolism, and susceptibility to infection (Round and Mazmanian 2009, Belkaid and Hand 2014). Animals capable of assessing the microbiome status of conspecifics could gain significant fitness advantages – for example, by avoiding parasitized or immunocompromised individuals, preferring mates with complementary or healthy microbiomes, or by selecting social partners that share beneficial microbial communities. Such behaviors have been documented or hypothesized in several species (Kavaliers, Choleris et al. 2005) (Arakawa, Cruz et al. 2011), but the sensory mechanisms supporting them have remained unclear. Our data suggest that the bile acid channel of the AOS may serve as one such mechanism, providing the animal with a real-time chemical readout of conspecific microbiome composition.

### Limitations and future directions

Several important questions remain. First, our experiments used germ-free mice as an extreme model of microbiome alteration. While this approach maximizes chemical contrast and provides clear proof of principle, naturally occurring microbiome variation is subtler. Whether VSN bile acid tuning can distinguish among conspecifics with more moderate differences – for instance, animals housed under different dietary or environmental conditions – remains to be determined. Second, while we have established the peripheral coding strategy at the level of the VNO, how this combinatorial information is integrated and potentially transformed in the accessory olfactory bulb and downstream brain circuits to generate behavioral outputs such as aversion or social investigation will be a critical next step. Third, the specific vomeronasal receptors mediating submicromolar taurine-conjugated bile acid detection remain unidentified. Future transcriptomic profiling of the functionally defined VSN clusters described here, combined with receptor deorphanization studies, will be essential for establishing the molecular basis of this sensitivity.

Together, our findings demonstrate that vomeronasal bile acid tuning is fundamentally mapped to the chemical characteristics of species, diet, and gut microbiome variations. The especially strong alignment between VSN tuning and microbiome-dependent bile acid patterns suggests that selective pressure from conspecific chemical evaluation may have been a potent driver of AOS bile acid receptor evolution. This interpretation is consistent with the emerging view that mammalian chemosensory systems coevolved with host-associated microbial communities (Elizabeth and Kevin 2011, Ezenwa and Williams 2014). More broadly, these studies demonstrate that a peripheral sensory population implements a stimulus whitened combinatorial code matched to the chemical output of gut physiology. This firmly establishes a robust across-animal chemosensory interface between the nervous system of an animal and gut characteristics of other vertebrates.

## MATERIALS AND METHODS

### Animals

All animal experiments were performed in accordance with the University Committee on Animal Resources, the Institutional Animal Care and Use Committee for the University of Rochester Medical Center, and follow the guidelines of the Guide for the Care and Use of Laboratory Animals. Physiological experiments were performed with OMP^tm4(cre)Mom/J^ knock-in mice (OMP-Cre mice; Jackson Laboratory Strain #006668)(Li, Ishii et al. 2004) mated with Gt(ROSA)^26Sortm96(CAG-GCaMP6s)Hze/J^ mice (Ai96 mice; Jackson Laboratory Strain #028866)(Madisen, Garner et al. 2015). Double-heterozygous (OMP-Cre^+/−^ and Ai96^+/−^) offspring from these mating pairs express the genetically encoded Ca^2+^ indicator GCaMP6s in VSNs, hereafter referred to as OMPxAi96 mice. 2–5 animals per cage were maintained on a 12-hour light/dark cycle and had *ad libitum* access to food and water. All experiments were performed on adult mice aged 8-16 weeks. The number, strain, and sex of animals used in each experiment are described in Results.

### Solutions and stimulus presentation

The full panel of bile acids tested is listed in Table 1 and included primary unconjugated, secondary unconjugated, taurine-conjugated, glycine-conjugated, and keto bile acid species. All chemicals were purchased from Steraloids Inc. (Newport, RI, USA), MedChemExpress (Monmouth Junction, NJ), or Cayman Chemical (Ann Arbor, MI). Stock solutions (20 mM) of all BAs were prepared in methanol and diluted to their final concentration in Ringer’s solution containing 115 mM NaCl, 5 mM KCl, 2 mM CaCl_2_, 2 mM MgCl_2_, 25 mM NaHCO_3_, 10 mM HEPES. and 10 mM glucose. Bile acid stimuli were diluted to a concentration range of 0.001 to 10 μM. Negative control (vehicle-only) stimuli consisted of Ringer’s solution containing 1:2000 methanol (the highest methanol concentration in any individual stimulus). Stimuli were applied for 15 seconds using an air pressure-operated reservoir via a 16-in-1 multi-barrel “perfusion pen” (AutoMate Scientific, Berkeley, CA, USA).

For behavioral experiments, 50 mg/ml DCA or TDCA was prepared in 1 ml female feces extract (FF). Exposure to TDCA, DCA, or FF was achieved depositing 50-100 μL of solution onto using cotton-tipped applicators. Fecal extracts from female mice were prepared by dissolving 5 g of feces in 50 ml of distilled water (dH_2_O). After the fecal mixture was stirred for two minutes, it was placed on ice and shaken continuously throughout the night. Following a 2-minute vortex on the second day, the feces mixture was centrifuged twice (10 min at 2400 × g at 4°C and 30 min at 2800 × g at 4°C) to homogenize it. The supernatant from two or more centrifuged samples was combined, passed through a 0.22 μm filter, placed in collection tubes, and used fresh or stored at -80 °C until just before use.

### Volumetric VNO Ca^2+^ imaging

Similar to previous studies (Turaga and Holy 2012, Wong, Nagel et al. 2018, Wong, Cao et al. 2020), VNOs were dissected and the vomeronasal epithelium was carefully removed under a dissection microscope (Leica Microsystems, Buffalo Grove, IL, USA) after deep isofluorane anesthesia and rapid decapitation. The vomeronasal epithelium was placed in a specially designed imaging chamber after being mounted on nitrocellulose paper (Thermo Fisher Scientific, Atlanta, GA, USA). A custom-made objective-coupled planar illumination (OCPI) microscope was used to perform volumetric Ca^2+^ imaging with previously reported refinements (Wong, Nagel et al. 2018, Wong, Cao et al. 2020). Fluorescence imaging experiments were supported by custom software that coordinated imaging and stimulus delivery through a randomized, interleaved stimulus delivery system (AutoMate Scientific). Each image stack encompassed approximately 700 μm laterally, 250 to 400 μm axially, and ∼150 μm in depth, with each image stack containing 51 frames, acquired once every 3 s (∼0.33 Hz). Five consecutive stacks (∼15 s) of each stimulus were delivered, with at least 10 stacks (≥30 s) elapsed between stimulus trials. Three or more full randomized, interleaved stimulus blocks were completed in each analyzed experiment.

### Data analysis of volumetric VNO Ca^2+^ imaging

Custom MATLAB software was used for data analysis similar to previous studies (Turaga and Holy 2012, Hammen, Turaga et al. 2014, Wong, Cao et al. 2020). Multiple rounds of rigid registration were used to align image stacks across time, supported by SimpleElastix software (Marstal, Berendsen et al. 2016). Following registration, regions of interest (ROIs) encompassing VSN somas were drawn in 3-dimension using a custom MATLAB user interface. A matrix of fluorescence intensity was produced by calculating the mean voxel intensity within each ROI for each of the 1200 image stacks in every experiment. To analyze the responses of VSN- encompassing ROIs, the mean voxel intensity in three successive pre-stimulus stacks was subtracted from the mean voxel intensity of three stacks during stimulus delivery, and the result was divided by the mean pre-stimulus intensity value to determine ΔF/F, the relative change in GCaMP6s intensity. We then calculated the across-trial mean ΔF/F for all ROIs and stimulus applications. In order to account for potential effects of valve switching, the across-trial ΔF/F for each ROI was compared to the Ringer’s control stimulus. ROIs exhibiting a positive ΔF/F response to any stimulus greater than 10%, with an across-trial and p-value less than 0.1 (Student’s two-tailed *t*-test) were included in subsequent analysis. Clustering was performed using density-based merging (DBM)(Walther, Zimmerman et al. 2009), part of the “runUMAP” package available in MATLAB. DBM parameters were selected based on manual inspection of results, and inspection of pairwise linear discriminant analysis (LDA), which was used to evaluate cluster separation via the discriminability index statistic (d′, see Fig. 2F). Satisfactory cluster separation was confirmed by a discriminability index score greater than 1.64 (when the z-score equivalent p-value is equal to 0.05).

### Mass spectrometry analysis

Reptile fecal samples were collected by members of the Division of Herpetology at the Dallas Zoo as part of routine cage maintenance and cleaning, and did not involve any alteration to standard animal housing routines. Fresh fecal samples were placed into plastic tubes and stored at -20 °C until collection by laboratory staff. Aqueous extracts from these samples were collected as noted above (Solutions and stimulus presentation), and aliquots were stored at -80 °C until use. Mass spectrometry analysis of bile acids in reptilian samples was performed similar to previous methods (Han, Liu et al. 2015). Briefly, verified monomolecular bile acid standards purchased from Steraloids, Santa Cruz Biotechnologies (Santa Cruz, CA, USA), Toronto Research Chemicals (Toronto, Ontario, Canada), or C/D/N Isotopes (Pointe-Claire, Quebec, Canada) and used as isotope-labeled standards. Samples were analyzed via ultrahigh performance liquid chromatography/multiple-reaction monitoring-mass spectrometry (UPLC-MRM-MS). This involved an Ultimate 3000 RSLC system (Dionex Inc., Amsterdam, The Netherlands) coupled to a 4000 QTRAP mass spectrometer (AB Sciex, Concord, ON, Canada) via a Turbo Ionspray electrospray ionization (ESI) source, which was operated in the negative ion mode. A BEH C18 (2.1 mm × 150 mm, 1.7 μm) UPLC column (Waters Inc., Milford, MA) was used for the gradient elution, with 0.01% formic acid in water (solvent A) and 0.01% formic acid in acetonitrile (solvent B) as the mobile phase. The collision energy for each group-specific MRM transition was the median of the collision energies for the same transition for all the isomeric bile acids in each group. Abundance measurements for each bile acid analyte were made for 3 replicates, and the mean abundance value utilized for subsequent analysis.

Germ-free mouse fecal extracts were obtained from adult male and female C57Bl6 mice housed in a gnotobiotic housing facility via University of Rochester’s Division of Comparative Medicine as part of routine cage maintenance. Aqueous extracts of germ-free fecal samples, as well as from C57Bl6 mice maintained in a standard/conventional mouse housing facility, were diluted in dH_2_O and submitted for analysis via Creative Proteomics (Shirley, NY, USA). Briefly, samples were analyzed via an AB SCIEX Qtrap 5500 mass spectrometer operated in negative ion mode, and connected to a Waters ACQUITY Ultra Performance Liquid Chromatographer. Samples were run over a Waters ACQUIRT UPLC BEH C18 column (2.1 mm × 100 mm, 1.7 μm) for the gradient elution, with 0.05% formic acid in water (solvent A) and 0.05% formic acid in acetonitrile (solvent B) as the mobile phase. Absolute abundance measurements (in ng/ml) for each bile acid analyte were generated for at least 2 technical replicates per sample, and averaged for subsequent analysis.

### Behavioral test setup

All behavioral tests were performed under dim red light during the dark cycle. Mice were solo housed for 10 days before the start of experiment. For nonsocial behavioral measurements of taurine-conjugated bile acid impact, we added TDCA (50 mg/ml) to female feces extracts (FFE), then dipping cotton-tipped applicators into TDCA+FFE or FFE alone as a control. Mice were habituated to the experimental cage for 10 minutes, followed by introduction of cotton swabs having either FF extract or FF+TDCA. Videos of animal behavior were taken for 10 min, and close inspection of each cotton swab manually evaluated by a blinded individual.

For social behavioral measurements of the impact of TDCA, mice were first maintained in groups of 3 for one week prior to behavioral testing. One day prior to the experiment, we collected fecal material from the cages and made aqueous fecal extracts as described above. Mice were acclimatized to behavior room one day prior to testing. On the day of experiment, two of the mice were separated from the test mice for 1 hour. Two mice were then anesthetized via ketamine-xylazine injection, and painted in the anogenital areas with either the pooled fecal extracts (PFE) or fecal extracts spiked with TDCA (50 mg/ml) at a volume of 200 μL. Test mice were allowed to habituate in the experimental cage for 10 min. The painted mice were then placed in two corners of the experimental cage, and videos of behavior recorded for 10 min. Close inspection of each anesthetized individual was manually evaluated by a blinded individual.

### Integration of neural and chemical bile acid data

We used two approaches to compare VSN bile acid tuning to mass spectrometry measurements. The first, used in the analyses presented in Figures 3 and 6, utilized neural datasets in which the original source of mass spectrometry data (*e.g.*, the reptilian fecal extracts in Fig. 3) was not used as a direct stimulus in the VSN imaging experiment. We will refer to this method of evaluating chemical/neural alignment as the “Indirect Method.” The second method, used in the analysis presented in Figure 7, utilizes neural datasets in which the source of mass spectrometry information was used as a direct VSN stimulus alongside representative monomolecular bile acid ligands at 1 μM. We will refer to this as the “Direct Method.”

### Data preprocessing

*Indirect Method:* Average raw mass spectrometry abundance measurements from reptilian feces were normalized on a per-sample basis to account for unknown and variable hydration states at collection. The maximum normalized abundance (1.0) for these samples was assigned the maximum value of 1000 arbitrary abundance units. Bile acid analytes that were undetectable in a sample were assigned a value of 0.0001 arbitrary abundance units. Normalized abundance measurements were log_10_ transformed, resulting in maximal values for each sample of +3 units, and minimum values (undetected analytes) of -4 units (see Figs. 2-3). For consistency, the same process was applied to germ-free and conventionally raised fecal extracts (Fig. 6). Normalized VSN responses and associated functional clusters were used as raw input for neural response correlates.

*Direct Method:* Chemical abundance data obtained via targeted mass spectrometry of germ-free (n=4 samples, 2 male, 2 female) and conventionally raised (n=2 samples, 1 male, 1 female) fecal extracts were preprocessed by scaling raw abundance values by a factor of 10^3^ and subsequently applying a log_10_ transformation to stabilize variance and accommodate the power-law distribution characteristic of metabolite abundances. Normalized VSN responses and associated functional clusters were used as raw input for neural response correlates (Fig. 7).

### Encoding models

#### Indirect Method

VSN bile acid tuning information was pooled across tissues and samples and functional clusters assigned as described above. Cluster assignments (n=20 for Fig. 3; n=19 for Fig. 6) were supplied as categorical variable labels to a MATLAB implementation of the “direct” linear discriminant analysis algorithm (Yu and Yang 2001), which identifies multidimensional eigenvectors (a.k.a. “discrimination axes”) that best separate functional VSN clusters from one another based on combinatorial response patterns. Similarly, normalized, log-transformed mass spectrometry abundance data were analyzed via the direct LDA algorithm. For reptile feces samples, we supplied the LDA algorithm with combined categorical variables of taxonomic suborder (4 in total) and diet (carnivore, vegetarian), resulting in 6 unique combinations and 5 resulting LDA eigenvectors (Fig. 3). For germ-free and conventionally raised mouse fecal samples, the same process involved just two categorical variables (germ-free and conventional), resulting in a single LDA eigenvector (Fig. 6). Pearson correlation analysis was performed pairwise on eigenvectors (similar to weight matrices) for pairs of neural- and mass spectrometry-derived LDA eigenvectors. Because the mathematical sign (positive or negative) of eigenvector weights are arbitrary, strong positive correlations and negative correlations equally indicate alignment of neural and mass spectrometry measurements. Correlation analyses between mass spectrometry-associated bile acid abundance patterns and VSN tuning (Fig. 3G, Fig. 6G-I) compared the maximum log_10_ normalized abundance values to the overall fraction of VSNs responses to each bile acid (those VSNs with ΔF/F values exceeding 5% and statistical p-values less than 0.01 compared to vehicle control, Student’s unpaired, two-tailed *t*-test, 3 or more stimulus repeats).

#### Direct Method

For each VSN, the normalized ΔF/F response to each of the eight individual bile acid stimuli (CA, LCA, CDCA, TCA, DCA, TDCA, TLCA, TCDCA) was used as a predictor variable, and the averaged ΔF/F response to CF or GF fecal extracts served as the response variable. Separate models were fit for the CF condition (averaged across CF male and CF female fecal extract stimuli) and the GF condition (averaged across GF male 1, GF female 1, GF male 2, and GF female 2 fecal extract stimuli). The population-level encoding model was formulated as:

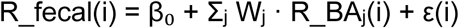

where R_fecal(i) is the averaged fecal extract response of neuron i, R_BAⱼ(i) is neuron i’s response to bile acid j (j = 1, …, 8), Wⱼ is the regression coefficient (encoding weight) for bile acid j, β₀ is the intercept, and ε(i) is the residual. Models were fit by ordinary least squares (OLS) regression across all VSNs using the MATLAB function *regress*, which also returned 95% confidence intervals for each weight and F-statistics for overall model significance.

### Weight–abundance correlation

#### Direct method

The encoding weight difference vector (ΔW = W_CF − W_GF) was computed element-wise across the eight bile acids, capturing the differential contribution of each bile acid to CF versus GF fecal extract encoding. Independently, the chemical abundance difference vector (ΔC = C_CF − C_GF) was computed from mass spectrometry data, where C_CF and C_GF represent the mean log₁₀-transformed abundances of each bile acid in conventional (n = 4) and germ-free (n = 8) fecal samples, respectively. The central hypothesis — that bile acids whose fecal abundances differ most between microbiome states should also show the largest differences in neural encoding weights — was tested by correlating ΔW with ΔC.

Pearson and Spearman rank correlations were computed between ΔW and ΔC. Because the sample size (N = 8 bile acids) limits the reliability of parametric inference, statistical significance was assessed by a two-tailed permutation test (10,000 iterations), in which the elements of ΔC were randomly shuffled and the Pearson correlation recomputed for each permutation. The permutation p-value was defined as the proportion of permuted correlations with absolute value greater than or equal to the observed correlation. A bootstrap procedure (10,000 iterations, resampling bile acids with replacement) was used to estimate the 95% confidence interval for the Pearson correlation coefficient. Individual weight–abundance correlations were also computed separately for the CF model (W_CF versus C_CF) and the GF model (W_GF versus C_GF), each with its own permutation test, to evaluate condition-specific alignment between neural and chemical information.

To control for the possibility that the ΔW–ΔC relationship was driven by overall response magnitude rather than discriminative tuning, a partial correlation was computed. Both ΔW and ΔC were residualized on their respective condition means (W_mean = (W_CF + W_GF)/2 and C_mean = (C_CF + C_GF)/2) by linear regression, and the Pearson correlation was computed between the residuals.

### Other analyses (Direct method only)

#### Per-stimulus robustness

To verify that the weight–abundance relationship was not an artifact of averaging across sex or biological replicates, the encoding model was fit separately for each of the six individual fecal extract stimuli (CF male, CF female, GF male 1, GF female 1, GF male 2, and GF female 2). For each per-stimulus weight vector, the correlation with ΔC was computed to assess consistency across stimuli.

#### Canonical correlation analysis

To evaluate the joint alignment of neural and chemical representations across both microbiome conditions simultaneously, canonical correlation analysis (CCA) was applied to the neural weight matrix [W_CF, W_GF] (8 × 2) and the chemical abundance matrix [C_CF, C_GF] (8 × 2). CCA identifies linear combinations of neural and chemical variables (canonical variates) that maximize cross-covariance, providing a multivariate test of neural–chemical alignment. Significance of the first canonical correlation was assessed via Wilks’ Λ using MATLAB’s *canoncorr* function.

#### Jackknife influence analysis

To assess whether the ΔW–ΔC correlation was dependent on any single bile acid, a leave-one-out jackknife analysis was performed. The Pearson correlation was recomputed eight times, each time omitting one bile acid, and the change in correlation (Δr) was recorded. The bile acid whose removal produced the largest |Δr| was identified as the most influential data point.

#### Sign concordance test

As a non-parametric, assumption-free test of directional agreement between neural and chemical discrimination, we evaluated sign concordance: for each bile acid, we determined whether ΔW and ΔC shared the same sign (both positive or both negative). Under the null hypothesis of no association, each bile acid has a 0.5 probability of sign agreement. Statistical significance was assessed using a one-sided binomial test.

#### Virtual knockout analysis

To quantify each bile acid’s contribution to the integrated neural–chemical discrimination signal, we computed a scalar discrimination score defined as the dot product of the encoding weight difference vector and the chemical abundance difference vector: S = ΔW · ΔC. This score represents the total alignment between neural tuning and chemical abundance along the microbiome discrimination axis. A leave-one-out virtual knockout was performed by setting each bile acid’s ΔC value to zero in turn and measuring the percentage reduction in |S| relative to the baseline. The signed contribution of each bile acid was computed as W_diff(j) × C_diff(j), capturing both the magnitude and direction of each bile acid’s role in discrimination. Cumulative impact was assessed by ranking bile acids by knockout impact and computing the running sum. A leave-two-out analysis was performed for all pairwise combinations to detect synergistic or redundant interactions, defined as cases in which the pairwise knockout impact deviated from the sum of the individual impacts by more than five percentage points.

### Software and reproducibility

All analyses were performed in MATLAB (MathWorks). Permutation and bootstrap procedures used 10,000 iterations unless otherwise stated. No corrections for multiple comparisons were applied across independent analyses (encoding model, discrimination axis, CCA), as these address distinct and complementary aspects of the same hypothesis. Bile acid abbreviations follow IUPAC nomenclature where applicable.

## Acknowledgments

We thank members of the Chemosensation and Social Learning Laboratory for helpful feedback. We especially thank Michael Mastrangelo for critical animal husbandry support. We thank the Dallas Zoo Department of Herpetology for providing reptile fecal samples (DZ Project S2019-2), and the laboratory of Dr. Felix Yarovinsky for providing gnotobiotic fecal samples.

## Author contributions

V.H. and J.P.M. designed the project, analyzed data, and wrote the manuscript. V.H. and W.M.W. performed microscopy experiments. V.H., J.W., M.M., and J.P.M. analyzed microscopy data. V.H. and L.R. performed behavior experiments. V.H. and J.P.M. analyzed behavioral data. J.W. and J.G.M collected reptile samples and conducted mass spectrometry experiments.

